# Trem2 deficiency does not worsen metabolic function in diet-induced obese mice

**DOI:** 10.1101/2022.06.13.495953

**Authors:** Nathan C. Winn, Elysa M. Wolf, Jamie N. Garcia, Alyssa H. Hasty

**Affiliations:** Department of Molecular Physiology and Biophysics, Vanderbilt University, Nashville, Tennessee, USA; VA Tennessee Valley Healthcare System, Nashville, Tennessee, USA

## Abstract

Triggering receptor expressed on myeloid cells 2 (Trem2) is highly expressed on myeloid cells and is involved in cellular lipid homeostasis and inflammatory processes. Trem2 deletion in mice (Trem2-/-) has been implicated in evoking adipose tissue dysfunction, but its role in worsening obesity-induced metabolic dysfunction is not resolved. Here we aimed to determine the causal role of Trem2 in regulating glucose homeostasis and insulin sensitivity in mice. Nine-week-old male and female littermate WT and Trem2-/- mice were fed low fat or high fat diet for 18 weeks and phenotyped for metabolic function. Diet-induced weight gain was similar between genotypes, irrespective of sex. Consistent with prior reports, we find that loss of Trem2 causes massive adipocyte hypertrophy and an attenuation in the lipid associated macrophage transcriptional response to obesity. In contrast to published data, we find that loss of Trem2 does not worsen metabolic function in obese mice. No differences in intraperitoneal glucose tolerance (ipGTT), oral GTT, or mixed meal substrate control, including postprandial glucose, non-esterified fatty acids, insulin, or triglycerides were found between WT and Trem2-/- animals. Similarly, no phenotypic differences existed when animals were challenged with stressors on metabolic demand (i.e., acute exercise or environmental temperature modulation) or when animals were challenged with a non-lethal dose of endotoxin. Collectively, we report a disassociation between adipose tissue remodeling caused by loss of Trem2 and whole-body metabolic homeostasis in obese mice. The complementary nature of experiments conducted gives credence to the conclusion that loss of Trem2 is unlikely to worsen glucose homeostasis in mice.

## INTRODUCTION

Triggering receptor expressed on myeloid cells 2 (Trem2) is a transmembrane receptor of the immunoglobulin superfamily that binds an array of ligands, including bacteria and polyanionic molecules (1), DNA (2), lipoproteins (3) and phospholipids (4). Trem2 is highly conserved between species and is primarily expressed on cells of the myeloid lineage. Trem2 is implicated in Alzheimer’s disease where its activation in microglia dampens the microglial inflammatory response [reviewed in (5)]. This has sparked interest in Trem2 as a therapeutic target in neurodegenerative diseases. Outside of the central nervous system, the role of Trem2 is less clear, but is being rapidly explored in a number of disease states including cancer and metabolic disease.

Recent data indicate that a macrophage population enriched in obese adipose tissue – termed lipid associated macrophages (LAMs) – is defined by high expression of *Trem2*/*TREM2* in both mice (6, 7) and humans (6). Mice devoid of Trem2 were reported to manifest with worsened glucose homeostasis and enhanced AT inflammation in the setting of diet-induced obesity (6, 8). This suggests that LAMs are lipid scavenging phagocytes that act as a buffer during adipose tissue expansion. Interestingly, overexpression of Trem2 also worsened glucose metabolism and adipose tissue dysfunction in mice (9), possibly due to ectopic expression of Trem2 in non-canonical cell types. Thus, the role by which Trem2 modulates glucose metabolism *in vivo* is unresolved. Given that anti-Trem2 therapies are being explored in early clinical trials, it becomes of the utmost importance to resolve whether Trem2 is an implicit regulator of macrophage function and by extension, glucose regulation. Moreover, Trem2 knockdown has been shown to decrease anti-inflammatory effects of exercise in rat microglia (10). Yet, whether Trem2 is an important responder and/or regulator of physical activity-induced metabolic demand outside the central nervous system is unknown.

In the present study, we aimed to i) provide clarity as to whether Trem2 is a molecular trigger influencing whole body glucose metabolism in lean and obese states, ii) determine whether Trem2 is necessary for peak aerobic capacity and/or endurance performance, and iii) determine whether loss of Trem2 heightens the inflammatory response to endotoxin. To this end, male and female mice were fed either low fat (LFD) or high fat (HFD) diet to induce obesity followed by extensive metabolic phenotyping. In contrast to prior reports, we find that Trem2 deficiency does not worsen metabolic function, exercise capacity, or the early phase inflammatory response to a bacterial pathogen, however, in concert with prior studies Trem2 is necessary for adipose tissue remodeling in obesity and to regulate the macrophage lipid transcriptional profile.

## METHODS

### Ethics Statement

All procedures were approved in advance and carried out in compliance with the Vanderbilt University Institutional Animal Care and Use Committee. Vanderbilt University is accredited by the Association for Assessment and Accreditation of Laboratory Animal Care International.

### Animals

C57BL/6J (#000664, JAX) and Trem2 knockout (#027197, JAX; donating investigator: Mike Sanser) mice were purchased from Jackson Laboratories and bred in our facility to generate male and female WT and Trem2-/- mice as littermate controls. Trem2-/- animals contain a 175 base pair deletion within exon 2 resulting in an early stop after amino acid 17. Genomic DNA was isolated from mouse kidney using a Qiagen DNAeasy Blood and Tissue kit. The deletion was confirmed using forward primer: TCAGGGAGTCAGTCATTAACCA and reverse primer: CAATAAGACCTGGCACAAGGA according to the protocol detailed by Jackson Laboratories (https://www.jax.org/Protocol?stockNumber=027197&protocolID=19636). A separate cohort of Trem2-/- mice and C57Bl/6J control mice (n=9/genotype) was purchased directly for several experiments. Data from littermates or non-littermates are indicated in figure legends. Mice were fed either LFD or HFD during interventions (**Fig. 1A**). Environmental housing temperature was either room temperature (RT, 22°C) or thermoneutral temperature (TN, 29°C) for all studies.

**Figure 1.**
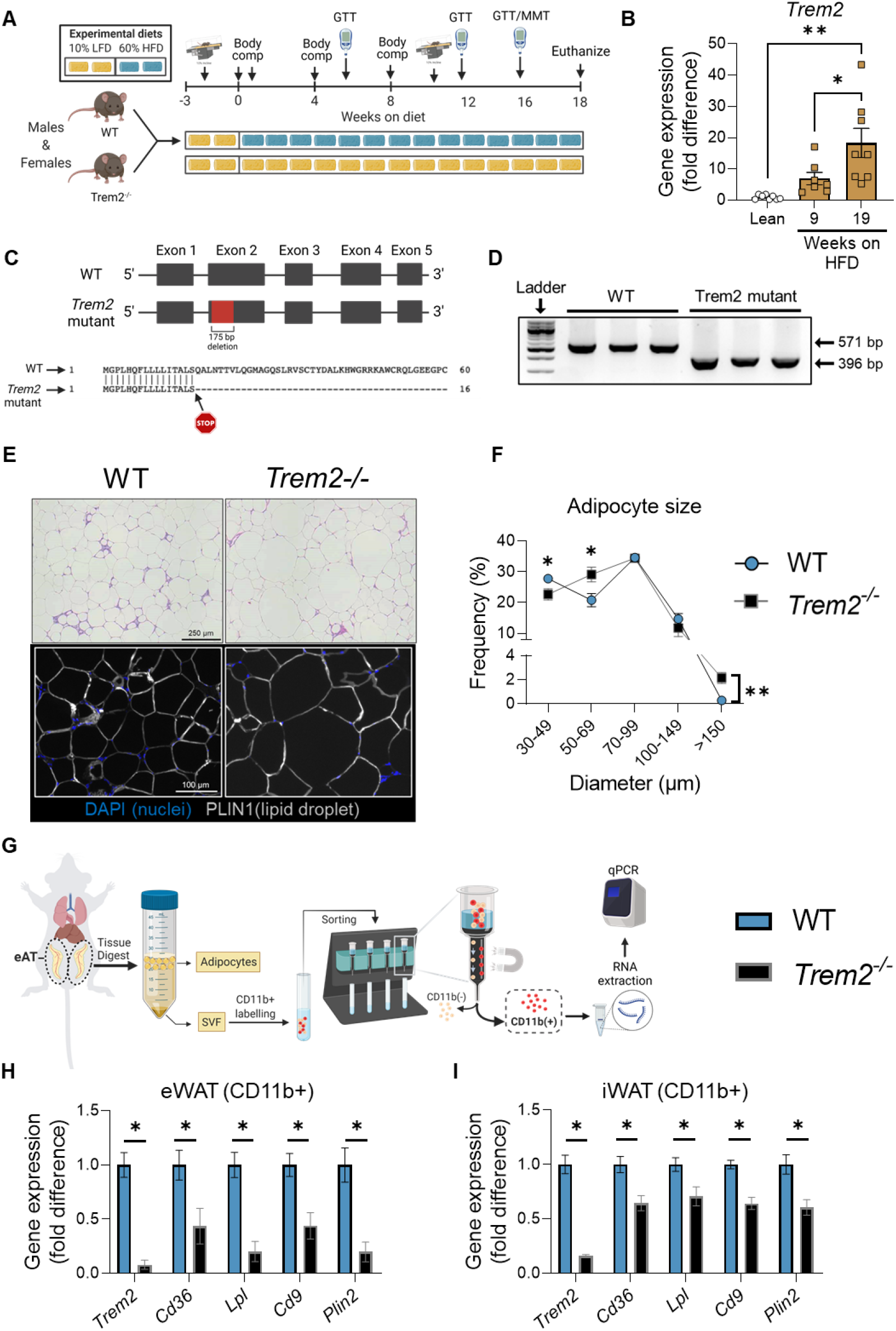
Trem2 deficiency enhances adipocyte hypertrophy and decreases transcriptional lipid signature in macrophages. **A**) Primary study design. Male and female WT and Trem2-/- littermate mice were fed LFD or HFD for up to 18 weeks with frequent metabolic phenotyping. **B**) *Trem2* qPCR of gonadal WAT after 9 weeks or 18 weeks of HFD feeding compared with lean controls. **C**) Representation of WT and Trem2-/- genomic DNA sequences. Trem2-/- animals (JAX#027197) are characterized by a 175 bp deletion within exon 2 resulting in an early stop within the cDNA after amino acid 17. **D**) PCR of WT and Trem2-/- gDNA confirming 175 bp deletion. **E**) Adipose tissue sections were generated and stained with H&E or DAPI/PLIN1 fluorescent antibodies to determine adipocyte size and morphological characteristics in 18 week HFD-fed mice. **F**) Adipocyte size distribution was computed using ImageJ, with sized adipocytes presented as diameter (μm). The frequency of a given bin size (e.g., diameter) was plotted on the y-axis. **G**) Epididymal and inguinal adipose tissue were isolated, digested, and separated by positive selection for CD11b+ macrophages in 18 week HFD-fed mice. **H**) qPCR was performed on CD11b+ fraction from epididymal adipose tissue and **I**) inguinal adipose tissue. One-way ANOVA was used to assess group differences for panel B with Tukey adjusted pairwise comparisons, (n=7-8/group). Independent t-tests were used to assess statistical differences between groups for panels F, H, and I, (n=7-9/group). Data are presented as mean ± SE. Graphics were generated using BioRender.com. Animal housing and experiments were conducted at room temperature (21-23°C). GTT, glucose tolerance test; MMT, mixed meal tolerance test; LFD, low fat diet, HFD, high fat diet; eWAT, epididymal adipose tissue; iWAT, inguinal adipose tissue. *p<0.05, **p<0.01

The temperature at which mice were housed is recorded in each figure legend. All *in vivo* procedures were performed at the temperature with which mice were acclimated – RT or TN, respectively.

### Body composition

Mouse body fat mass and fat free mass (FFM) were measured by a nuclear magnetic resonance whole body composition analyzer (Bruker Minispec).

### Glucose tolerance testing

After a 5 h or 15 h fast basal blood glucose levels were measured (-10 min) followed by intraperitoneal (i.p.) glucose bolus (2.0 g dextrose/kg FFM) or oral gavage (3.0 g/kg FFM or 3.5 g/kg FFM). Blood glucose was sampled via tail cut at 15, 30, 45, 60, 90,120, and 150 minutes after injection using a hand-held glucometer (Bayer Contour Next EZ meter). Glucose area under curve from baseline was calculated using the trapezoidal rule.

### Meal tolerance testing & arterial catheterization

After 15 weeks of HFD feeding, a subgroup of male and female mice underwent carotid artery catheterization. The Vanderbilt Mouse Metabolic Phenotyping Center (VMMPC) performed surgeries. Detailed procedures are available at the following website (www.vmmpc.org). One week after surgery, mice were challenged with a mixed meal test (EnsurePlus®, 1.48 kcal/mL; dose = 12 kcal/kg body weight). After a 5 h fast, a baseline blood sample was obtained, after which animals were administered the liquid meal via oral gavage. Blood was collected at -10, 10, 60, 120, and 180 min. Plasma glucose, insulin, NEFA, and triglycerides were determined.

### Lipopolysaccharide (LPS) challenge

After 18 weeks of LFD diet feeding, 28-week-old WT and Trem2-/- mice were challenged with an i.p. bolus of LPS (2 mg/kg body weight; L4391, lot#0000130083, MilliporeSigma). Four hours after injection, mice were euthanized, blood was collected, and tissues (epididymal adipose tissue, liver, and spleen) were harvested and flash frozen. TNFα and IL6 cytokines were determined in plasma and tissue lysates by ELISA (BioLegend, IL6, #431304; TNFα, #430904). Cytokine expression in tissue lysates are expressed as pg/mg protein concentration.

### Exercise testing

Prior to exercise testing animals were acclimated (10-15 min in duration) on 2-3 occasions to a belt treadmill (Columbus Instruments, Columbus, OH). Mice were placed on the treadmill for 10 min without belt movement (0 m/min). Thereafter, the belt was turned on and mice ran ‘sub-maximally’ at 10 m/min (5% grade) for 10 min. The 2^nd^ acclimation session consisted of 15 min of exercise at 10 m/min (15% grade). If a 3^rd^ acclimation session was performed it was a replication of the 2^nd^ session. The exercise session was at least 48 h after the last acclimation bout.

#### Exercise tolerance

The speed of the treadmill was set at 10 m/min with 10 degrees incline. Treadmill speed was increase by 4 m/min every 3 minutes until exhaustion. An electrical grid is located at the end of the end of the treadmill. The electrical stimulus was set at 163 volts with a stimulus rate of 4 pulses/sec. The amperage was set at 1.0 milliamps. Animals were determined to be exhausted, if they refused to continue running and remained on the grid for more than 5 seconds, upon which they were immediately removed.

#### Exercise endurance

The speed of the treadmill was set to 50-55% of a mouse’s maximal running speed (obtained from exercise tolerance test). Following a brief warm-up, mice ran at a constant speed for 90 min. If mice exceeded 90 min running time, the speed was increased 10-20% until mice were fatigued. In another group of mice, a fixed speed of 15 m/min (10% grade) until exhaustion (approximately 90 min) was performed in lean WT mice to establish whether Trem2 expression is responsive to an acute bout of exercise in epididymal and inguinal adipose tissue (Fig. S1A).

#### Voluntary physical activity

A voluntary wheel running test was conducted over a 48h period. Mice were single housed during the observation period and recombined with cage-mates after the test. Mice were transferred to a cage containing a running wheel that was physically locked for a 24-h habituation period. The following day the wheels were unlocked and animals were allowed to run voluntarily. Wheel distance and average speed was recorded using a computer odometer (Artyea KINGMAS LCD Bicycle Bike Computer Odometer Speedometer Sd-548b).

### Immunohistochemistry and Fluorescence Microscopy

Epididymal adipose tissue sections were fixed in 4% PFA for three h and transferred to 70% ethanol at 4°C. Paraffin embedded sections were stained with H&E or immunolabeled with PLIN1 (Cell Signaling Technology, #3470, 1:1000 dilution), goat anti-rabbit IgG secondary (Abcam, Alexa Fluor® 647, #ab150079, 1:1000 dilution) and DAPI (Thermofischer, #62247, 1:4000 dilution). Slides were imaged using either a 10X objective on a Leica DMI8 widefield microscope and captured with a Leica DFC9000GT camera. Adipocyte sizing was performed as previously described (7).

### Bone marrow-derived macrophages (BMDM)

Tibia bones were removed and cleaned of tissue. Ends of bones were cut and flushed with RPMI (1%FBS). Collected marrow was passed through a 25-gauge needle several times slowly to create a single cell suspension. After centrifugation and decantation, the cell pellet was treated with ACK lysis buffer to remove red blood cells. Reaction was neutralized by dilution with 1% FBS in PBS. Cells were centrifuged, and supernatant decanted; cell pellet was resuspended in BMDM differentiation media (DMEM with 10% L929 conditioned media, 10% fetal bovine serum (FBS), 2 mM glutamine, 10 U/mL penicillin/streptomycin) and distributed into tissue culture plates for growth. Differentiation media was replaced on day 5. Following seven days of differentiation, cells were washed twice with pre-warmed 1x PBS and activated with LPS (10 ng/mL) and interferon gamma (IFNγ, 100 ng/mL) generating the following conditions: no inflammation (1x PBS), acute inflammation (LPS+IFNγ x 24 h), and inflammatory resolution (LPS+IFNγ x 24h followed by 48 h washout). After activation, cells were washed twice with pre-warmed 1x PBS, and isolated for RNA extraction.

### Adipose tissue macrophage isolation

Mice were euthanized by isoflurane overdose and cervical dislocation followed by perfusion with 20 ml PBS through the left ventricle. Epididymal and inguinal adipose tissue depots were excised, minced, and digested in 6 ml of 2-mg/mL type II collagenase (Worthington # LS004177) for 30 min at 37°C. Digested adipose tissue was then vortexed, filtered through 100 μm filters, and lysed with ACK buffer, and filtered through 35 μm filters as previously described (11). The stromal vascular fraction was incubated with anti-CD11b microbeads (Miltenyi) and positive selection was performed using gravity operated magnetic columns. The positive fraction (CD11b+ cells) was collected, washed 3 x with 1X PBS 1% FBS, and reconstituted in RLT lysis buffer (Qiagen #1015750) supplemented with 1% betamercaptoethanol. Samples were flash frozen and stored at -80°C until analysis.

### RNA isolation, cDNA synthesis, and real-time RT-PCR

Tissues or cells were homogenized and lysed in RLT buffer (Qiagen #1015750) with 1% betamercaptoethanol. Equal parts isopropanol and RNA-containing sample were transferred to a clean 1.5 mL tube. Purified RNA was reverse transcribed by iScript RT (Bio-Rad, Hercules, CA) into cDNA. Differences in relative gene expression were quantified using FAM-conjugated TaqMan Gene Expression Assay (Life Technologies). PCR reactions were performed in duplicate under thermal conditions as follows: 95°C for 10 min, followed by 40 cycles of 95°C for 15 s, and 60°C for 45 s. Data were normalized to *Gapdh*. mRNA expression values were calculated by the -ΔΔ cT method.

### Statistical Analyses

Student’s *t*-tests were run for between group comparisons. In experiments that contained more than two groups, one-way analysis of variance (ANOVA) or two-way ANOVA models were conducted with pairwise comparisons using Tukey or Sidak correction. Brown-Forsythe correction was applied to groups with unequal variance. Data are presented as mean ± standard error (SE). An adjusted *p* value of <0.05 was used to determine significance.

## RESULTS

### Trem2 deficiency enhances adipocyte hypertrophy and decreases transcriptional lipid signature in macrophages

Lipid associated macrophages are highly abundant in states of adipose tissue expansion and are defined by expression of Trem2 in humans and rodents (6-8). Indeed, a linear increase in *Trem2* expression manifests in response to diet induced obesity in adipose tissue (**Fig. 1A-B**). To confirm the 175 bp deletion in Trem2-/- mice, we performed PCR on genomic DNA isolated from the kidney of both WT and Trem2-/- mice. (**Fig. 1C-D**). Adipose tissue microscopy demonstrated that Trem2 null mice did not have greater mean adipocyte size (WT, 70±1.5 μm versus Trem2-/-, 72±2 μm, p=0.4) but the proportion of extremely large (>150μm diameter) adipocytes were significantly greater relative to body weight matched controls (**Fig. 1E-F**). This is similar to what was previously observed in other Trem2-/- models (6, 8). Gene expression from sorted myeloid cells (CD11b+) in epididymal and inguinal adipose tissue using positive selection (**Fig. 1G**) showed an 8-fold knockdown of *Trem2* compared with WT animals.

Similarly, genes associated with LAMs (*Cd36, Lpl, Cd9*, and *Plin2*) were decreased in Trem2 knockout cells providing additional support for loss of Trem2 function (**Fig. 1H-I**). Together, these data confirm the predicted phenotype from loss of Trem2 in adipose tissue macrophages.

### Trem2 is associated with inflammatory resolution

To determine whether loss of Trem2 potentiates systemic and tissue inflammation, mice were challenged with a sublethal dose of LPS (2 mg/kg BW; i.p.) and euthanized after 4 h (**Fig. 2A**). Body weight and blood glucose levels were not different between genotypes at baseline (**Fig. 2B-C**). As expected, after LPS exposure, blood glucose decreased but was not different between Trem2-/- and WT mice. Plasma IL-6 and TNFα were determined at baseline and 4 h post LPS exposure. In addition, IL-6 and TNFα were determined in epididymal adipose tissue, liver, and spleen. Loss of Trem2 did not augment tissue inflammation or alter plasma IL-6 or TNFα relative to WT controls (**Fig. 2D-G**). These data suggest that loss of Trem2 does not worsen early phase inflammatory response to a bacterial pathogen. Prior data suggest that Trem2-/- mice have accelerated death rates compared to WT controls in response to a lethal bacterial load (12). Given that no differences in the acute inflammatory response were found in this study, we posited that Trem2 may be important during inflammatory resolution. To test this relationship, BMDMs were generated from WT animals and challenged with inflammatory stimuli aimed at mimicking acute inflammatory phase and inflammatory resolution phase, respectively (**Fig. 2H**). *Trem2* expression was suppressed during acute inflammation (24 hours in vitro) and increased above basal during the resolution phase (**Fig. 2I**). An inflammatory score was computed as the sum of *Tnf, Cd38*, and *Nos2* and plotted against *Trem2* expression at each respective activation condition (i.e. no inflammation, acute inflammation, and resolution). In this plot, it is clear that *Trem2* is down-regulated with acute inflammation yet is upregulated during the inflammatory resolution phase (**Fig. 2J**).

**Figure 2.**
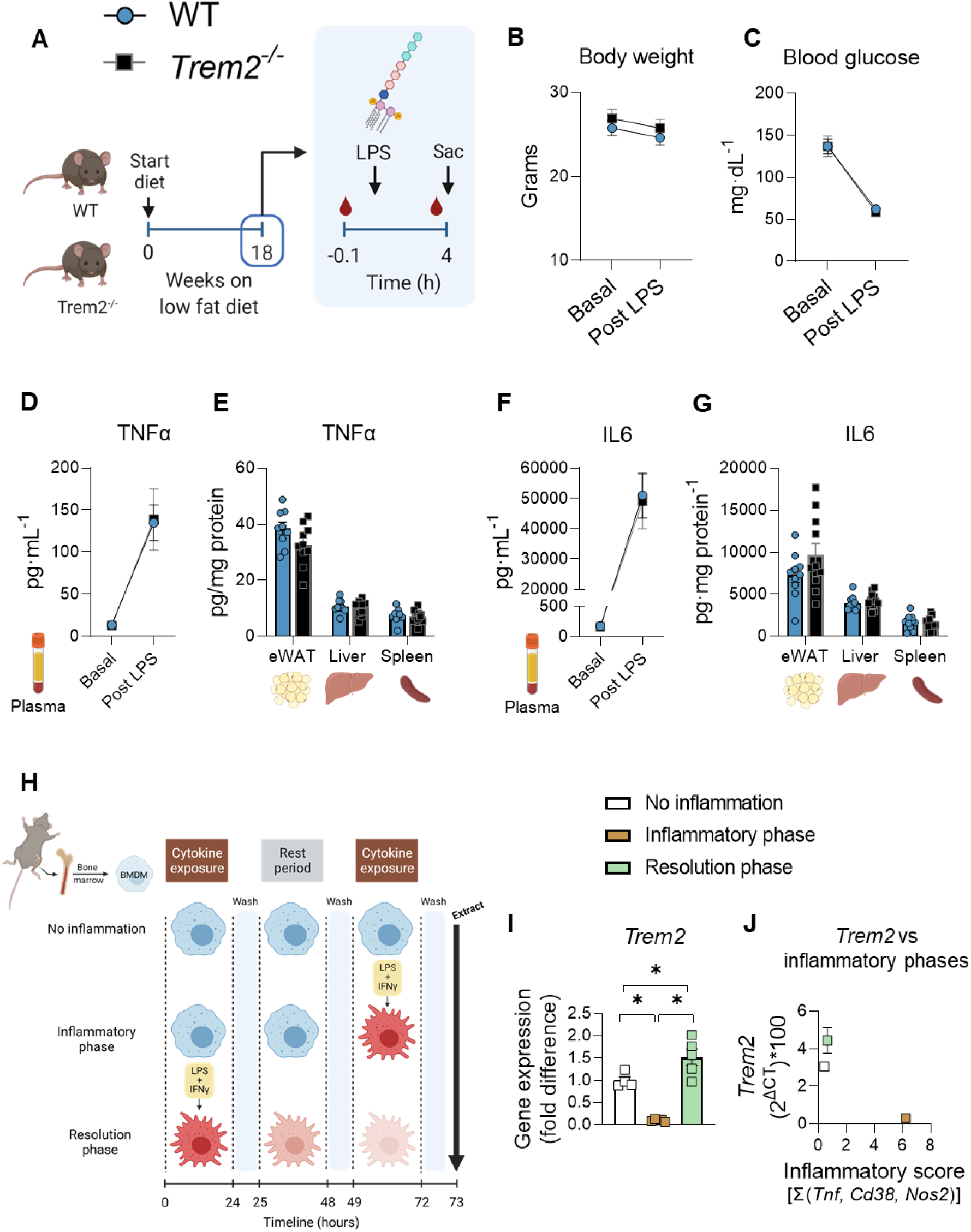
Trem2 is associated with inflammatory resolution. **A**) After 18 weeks on LFD, WT and Trem2-/- mice were challenged with a single bolus of lipopolysaccharide (LPS, 2 mg/kg body weight). Blood was collected at baseline and 4 h after LPS injection. **B**) body mass and **C**) blood glucose were monitored before and after LPS exposure. TNFα was determined in **D**) plasma and **E**) tissues.IL6 was determined in **F**) plasma and **G**) tissues. **H**) Bone marrow derived macrophages (BMDM)s were generated in vitro and exposed to control (no inflammation), acute activation with LPS+IFNγ (inflammatory phase), or prior activation with LPS+IFNγ (inflammatory resolution). BMDMs were extracted and mRNA expression was quantified. **I**) Trem2 mRNA expression in BMDMs and **J**) the relationship between Trem2 expression and inflammatory score during no inflammation, inflammatory phase, and inflammatory resolution phase. The inflammatory score was computed as the sum of three inflammatory genes (*Tnf, Cd38, and Nos2*). Independent t-tests were used to assess statistical differences between groups for panels E and G (n=10-11/group). Two-way repeated measures ANOVA with time and genotype as factors were conducted for panels B, C, D, and F (n=10-11/group). Multiple comparisons were assessed using Tukey *post hoc* testing. One-way ANOVA was used to assess group differences for panel I with Tukey adjusted pairwise comparisons (n=5/group). Blood was collected via retroorbital bleeding under mild anesthesia. Data are presented as mean ± SE. Graphics were generated using BioRender.com. Littermate controls were used for these experiments. Animal housing and experiments were conducted at room temperature (21-23°C). LPS, lipopolysaccharide; TNFα, tumor necrosis factor alpha; IL6, interleukin-like 6; BMDM, bone marrow-derived macrophage. *p<0.05

### Loss of Trem2 does not impair glucose homeostasis

To determine the metabolic consequences of Trem2 ablation, male and female WT and Trem2-/- mice were fed LFD or HFD diet for 18 weeks. No differences in body weight gain, feeding efficiency, or body fat mass were found between genotypes regardless of diet (**Fig. 3A-F**). Glucose tolerance was assessed at 12 and 16 weeks on diet. Blood glucose clearance was delayed in HFD versus LFD animals in both males and females, but Trem2 deletion did not worsen glucose tolerance after 12 weeks or 16 weeks of diet feeding, respectively (**Fig. 3G-J**). Given the discrepancy in glucose tolerance between the present study and previous reports (6, 8), we examined more closely the assessment of glucose tolerance in previous publications.

**Figure 3.**
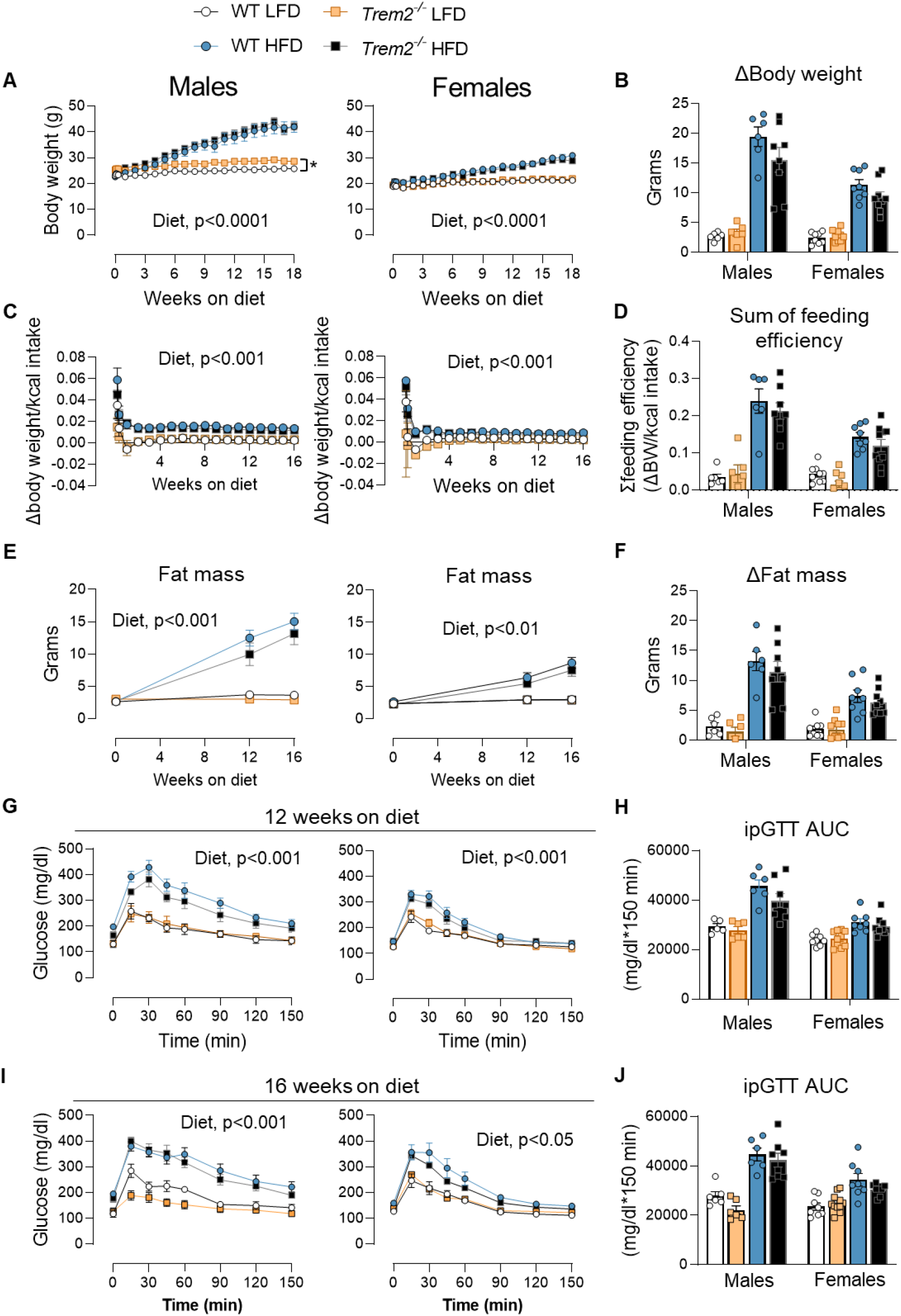
Loss of Trem2 does not impair glucose homeostasis. Male and female WT and Trem2-/- mice were fed LFD or HFD diet for up to 18 weeks. **A**) Weekly body mass was recorded throughout the feeding paradigm and **B**) the change in body weight from week 0 to week 18 was computed. **C**) Feeding efficiency was calculated throughout the study (feeding efficiency = change in weekly body weight per energy intake) and **D**) the sum of weekly feeding effcieincy was computed. **E**) Fat mass was measured over time via EchoMRI in addition to **F**) the fat mass delta from baseline. ipGTT was performed following a 5h fast **G&H**) at 12 weeks and **I&J**) 16 weeks of diet intervention (n=6-8/group). Two-way repeated measures ANOVA with time and group (diet and genotype) as factors were conducted for panels A, C, E, G and I (n=6-8/condition). Multiple comparisons were assessed using Tukey *post hoc* testing. Two-way ANOVA was used to assess group differences for panel B, D, F, H, and J with Tukey adjusted pairwise comparisons (n=6-8/group). Between sex comparisons were not included in the model, but male and female data are placed on the same scale. Blood was collected from the tail for these experiments. Graphics were generated using BioRender.com. All mice used in these experiments were littermates. Animal housing and experiments were conducted at room temperature (21-23°C). Data are presented as mean ± SE. LFD, low fat diet; HFD, high fat diet; ipGTT, intraperitoneal glucose tolerance test; AUC, area under the curve. *p<0.05

Each prior study fasted mice overnight (12-16 hours). Overnight fasting induces a catabolic state and enhances insulin action and glucose clearance in mice (13-16). Thus, to determine whether this catabolic state was necessary to unmask loss of Trem2-induced glucose intolerance, we overnight fasted a group of male mice fed HFD for 16 weeks and performed an ipGTT. First, glucose tolerance was determined at 6 weeks and 12 weeks of HFD feeding in the same cohort of mice to establish the change in glycemic control over time in Trem2-/- after short-term fasting (5h). No differences between genotypes were noted (**Fig. 4A**). ipGTT was conducted at 16 weeks of HFD feeding preceded by a 15h fast. Despite replicating fasting conditions from previous studies, we did not find worsened glucose tolerance in diet-induced obese Trem2-/- mice (**Fig. 4A**). There was a trend (GTT AUC, p=0.078) for greater glucose clearance in Trem2-/- than WT control mice. Fasting insulin and NEFA concentrations were not different between genotypes after 6 weeks, 12 weeks, or 16 weeks of diet feeding (**Fig. 4B-C**). Since the route of glucose administration can affect kinetics of glucose appearance and disappearance from circulation (17), and given that a previous publication found impaired glucose tolerance in Trem2-/- mice after oral gavage (8), we conducted an oral glucose tolerance test (OGTT) in obese male and female mice after 12 weeks of HFD feeding. Neither obese male nor female Trem2-/- mice had differences in blood glucose responses compared to WT controls (**Fig. 4D**). Fasting triglycerides and cholesterol were also similar between genotypes in both males and females (**Fig. 4E**).

**Figure 4.**
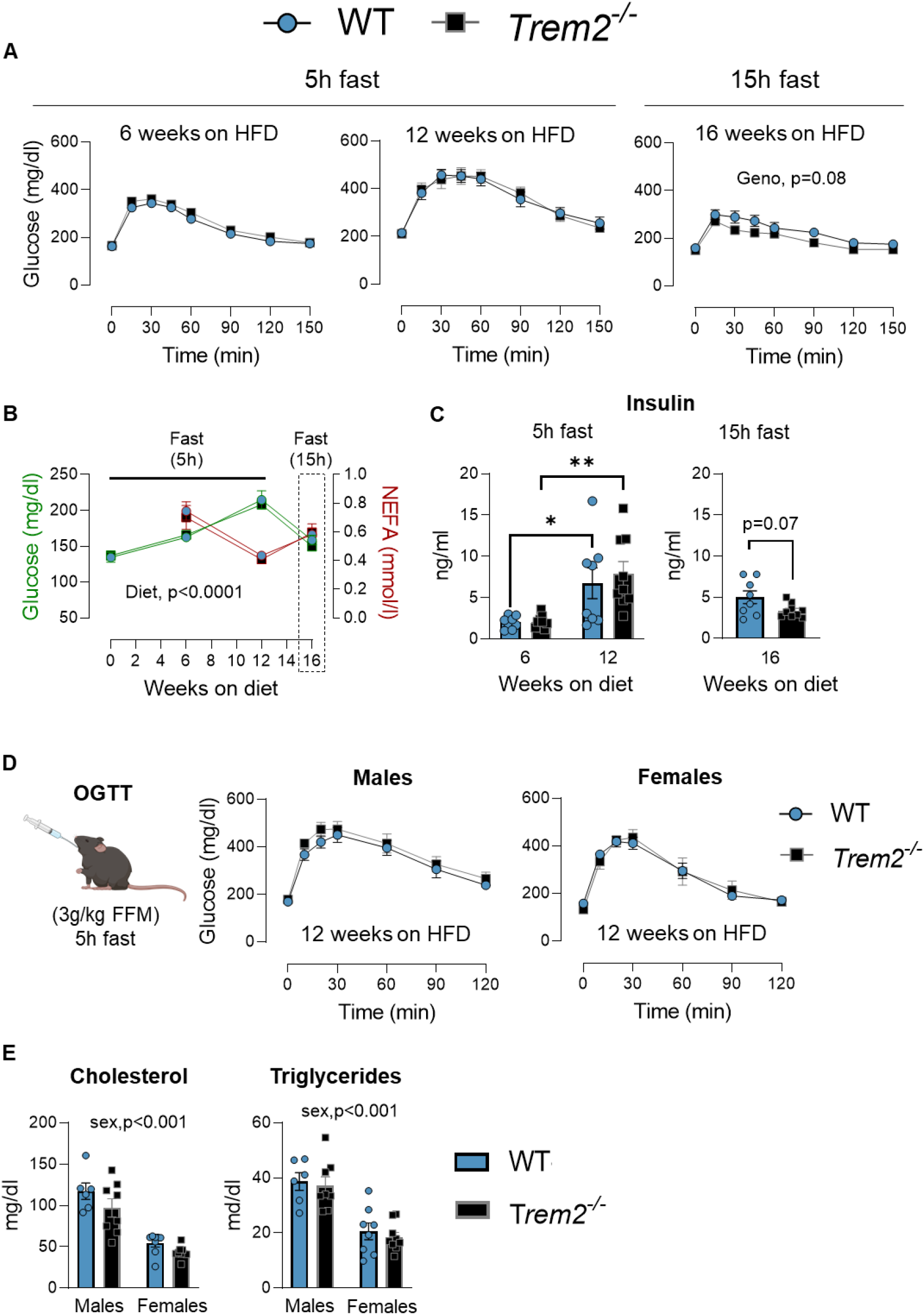
Neither extended fasting nor oral glucose challenge worsens glucose tolerance in Trem2-/- mice. Male Trem2-/- and WT mice were fed HFD diet for 16 weeks. **A**) ipGTT was performed after 6 and 12 weeks on HFD diet after a 5h fast. Another ipGTT was conducted at 16 weeks on HFD diet after a 15 h fast. **B**) Fasting glucose, NEFA, and **C**) plasma insulin concentrations were measured throughout diet feeding after either a 5 h of 15 h fast. **D**) oral glucose tolerance test (OGTT) was performed in male and female mice after 12 weeks on HFD diet and a 5 h fast. **E**) Fasting cholesterol and triglycerides were determined 12 week HFD fed males and females. Two way repeated measures ANOVA with time and genotype as factors were conducted for panels A, B, C, and D (n=6-9/condition). Multiple comparisons were assessed using Tukey *post hoc* testing. Two-way ANOVA with genotype and sex as factors was conducted for panel E with Tukey adjusted pairwise comparisons (n=6-8/group). In panels A, B, and C Trem2-/- mice were purchased directly from JAX and age-matched C57BL/6J mice were used as controls. Blood was collected from the tail for these experiments. Animal housing and experiments were conducted at room temperature (21-23°C). In panels D and E, mice are littermates. Graphics were generated using BioRender.com. Data are presented as mean ± SE. HFD< high fat diet; OGTT, oral glucose tolerance test; FFM, fat free mass. *p<0.05, **p<0.01

### Trem2 ablation augments feeding efficiency at thermoneutrality but does not decrease glucose tolerance

The ambient temperature at which an animal is housed exerts a profound effect on energy metabolism and glucose utilization. Indeed, energy expenditure of mice housed at RT is 50-100% higher than animals housed at TN (18, 19). Thus, we determined whether glucose tolerance in thermoneutral conditions was worse between Trem2-/- and WT mice. Mice of each genotype were housed at RT (22°C) or TN (29°C) while consuming LFD diet for three weeks and glucose tolerance was assessed (**Fig. 5A**). Body weights were similar between genotypes at the start of temperature exposure and did not differ over the 3-week period (**Fig. 5B**). Trem2-/- mice had higher feeding efficiency than WT mice at thermoneutrality while consuming LFD diet (**Fig. 5C-D**), suggesting higher fat balance, whereas no genotypic differences in energy homeostasis were found at standard RT. Oral glucose tolerance was not different between genotypes (**Fig. 5F**). Postprandial insulin was higher in animals housed under TN conditions, but no differences between Trem2-/- and WT mice were observed (**Fig. 5G**).

**Figure 5.**
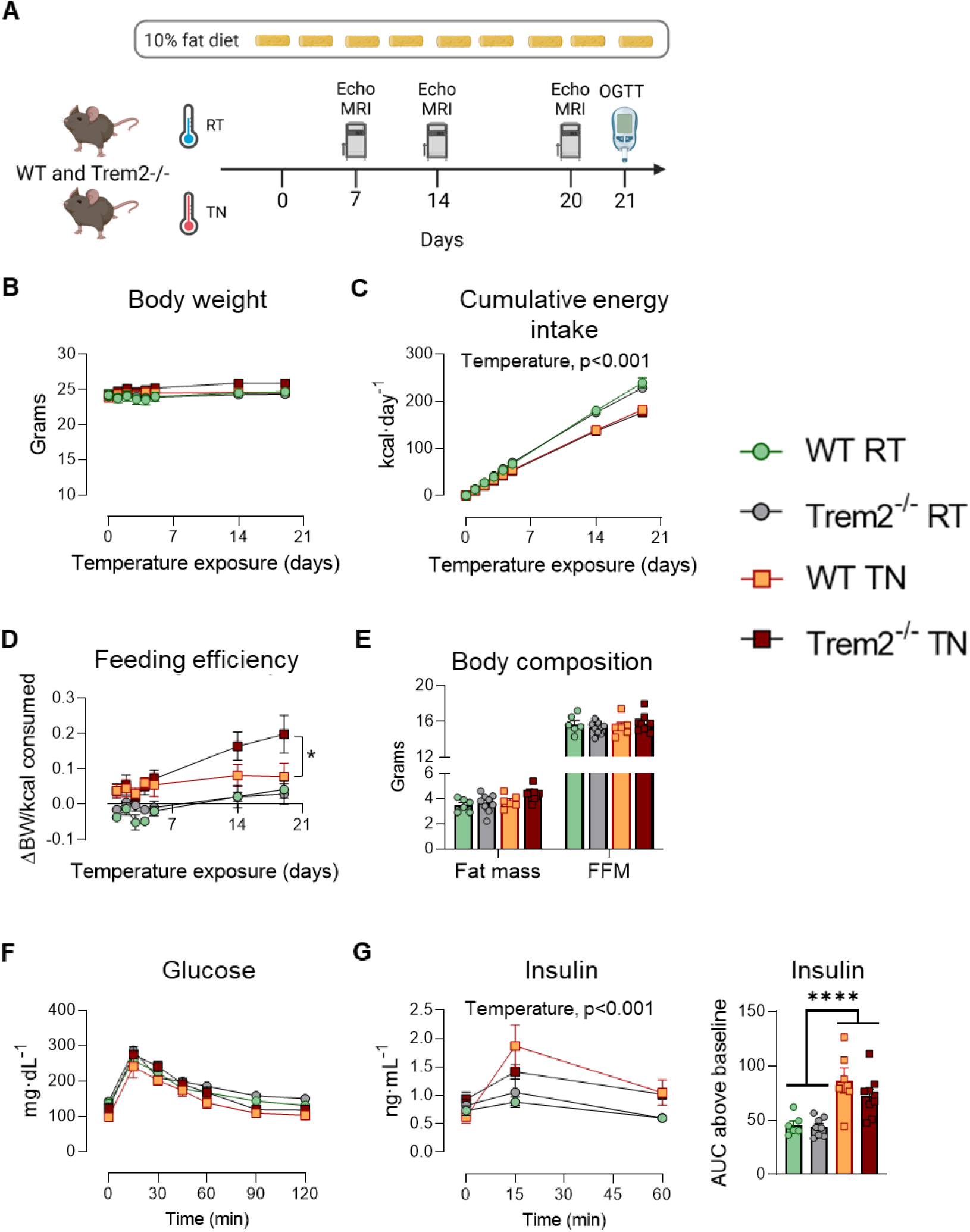
Trem2 ablation augments feeding efficiency at thermoneutrality but does not decrease glucose tolerance. **A**) Male Trem2-/- and WT mice were fed LFD diet for three weeks at standard room temperature (RT) or thermoneutrality (TN). **B**) Body weight, **C**) cumulative energy intake, and **D**) feeding efficiency were recorded frequently over time. After 21 days of temperature exposure **E**) body composition and **F**) OGTT were performed. **G**) Plasma insulin levels and insulin AUC were assessed. Two-way repeated measures ANOVA with time and group (genotype and housing temperature) as factors were conducted for panels B, C, D, F, and G (n=6-9/condition). Multiple comparisons were assessed using Tukey *post hoc* testing. Two-way ANOVA with genotype and housing temperature as factors was conducted for panel E and G (insulin AUC), with Tukey adjusted pairwise comparisons (n=6-9/group). All mice used in these experiments were littermates. Blood was collected from the tail for these experiments. Animal housing and experiments were conducted at the temperature with which mice were acclimated (RT, 22°C or TN, 29°C). Graphics were generated using BioRender.com. Data are presented as mean ± SE. RT, room temperature; TN, thermoneutral. *p<0.05, ****p<0.0001

### Mixed meal tolerance is not worsened by loss of Trem2

In a subset of obese mice, we next tested whether mixed meal substrate clearance was impaired by Trem2 deficiency. Mice were surgically implanted with carotid artery catheters and gavaged orally with a liquid meal (12 kcal/kg body weight) one week after surgeries (**Fig. 6A**). Neither fasting glucose nor postprandial glucose were different between genotypes (**Fig. 6B**). Insulin concentrations in the fasting and fed states did not differ between groups (**Fig. 6C**).

**Figure 6.**
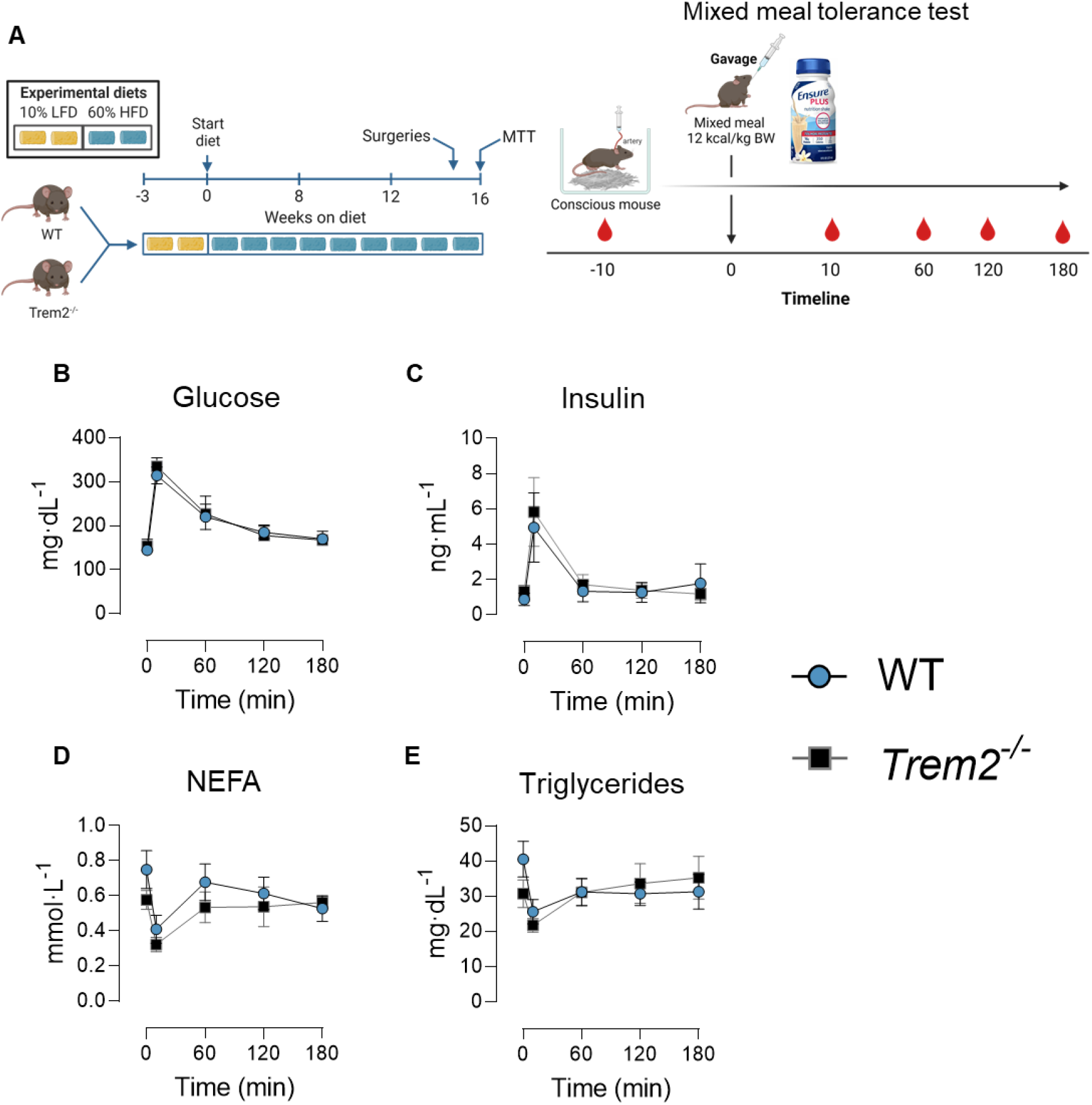
Mixed meal tolerance is not worsened by loss of Trem2. **A**) After 15 weeks of HFD feeding mice were surgically implanted with carotid artery catheters. Following one week of recovery, 5 h-fasted animals were challenged with an oral mixed meal (Ensure® 12 kcal/kg body weight). B) blood glucose, C) plasma insulin, D) plasma NEFA, and E) plasma triglycerides were determined in fasted and postprandial states. Two-way repeated measures ANOVA with time and genotype as factors were conducted for panels B-E (n=6-8/condition). Multiple comparisons were assessed using Tukey *post hoc* testing. All mice used in these experiments were littermates. Blood was collected from the carotid artery in an unrestrained and non-anesthetized animal. Animal housing and experiments were conducted at room temperature (21-23°C). Graphics were generated using BioRender.com. Data are presented as mean ± SE. NEFA, non-esterified fatty acids; MTT, mixed meal tolerance test.

Fasting NEFA and fasting triglycerides were slightly higher in WT animals but this did not reach statistical significance. Postprandial NEFA and triglycerides were similar between WT and Trem2-/- mice (**Fig. 6D-E**).

### Trem2 deletion does not limit acute exercise performance

Exercise is used clinically to uncover symptoms so as to facilitate disease diagnosis. It is possible that an increase in metabolic demand evoked by exercise unmasks a phenotype in Trem2-/- mice. Thus, we determined whether Trem2-/- mice were exercise intolerant relative to WT controls. Following treadmill acclimation sessions, mice underwent an exercise tolerance test [peak exercise test, (**Fig. 7A**)]. No differences in exercise tolerance were noted between genotypes (**Fig. 7B**). Since Trem2 expression is increased with obesity (**Fig. 1B**), we determined whether endurance capacity [low-moderate exercise test, **Fig. 7C**)] was limited in obese Trem2-/- mice. Nine weeks of HFD feeding decreased exercise endurance – assessed via time to exhaustion – but there were no differences between WT and Trem2-/- mice (**Fig. 7D**). Next, we determined whether voluntary running behavior was impacted by Trem2 deletion by giving mice access to running wheels (**Fig. 7E**). Mice were transferred to cages containing a running wheel for a total of 2 days. The first 24 h the running wheel was locked, and served as an acclimation period to the new environment. The next day the wheel was unlocked and mice were allowed to run voluntarily. Run distance, food intake, and body weight were recorded during the 48 h test (**Fig. 7F-G**). Neither voluntary run distance nor average wheel speed were different between groups (**Fig. 7H**). Recent data suggest that in rat microglia, Trem2 is involved in anti-inflammatory responses evoked by exercise. Given the absence of differences between genotypes in response to physical activity stressors, we determined whether Trem2 expression in epididymal and inguinal adipose tissue is responsive to acute exercise (**Fig. S1A**). *Trem2* expression was not significantly altered after a single endurance bout of treadmill exercise (mice were euthanized 4 h post exercise) (**Fig. S2D**). Notably, acute exercise exerted the anticipated response to lower plasma glucose and increase NEFA (**Fig. S2E-F**), thus the lack of Trem2 response in adipose tissue is unlikely related to inadequate lipolysis. Collectively, these data suggest that Trem2 is dispensable for acute exercise performance in lean and obese mice and does not influence voluntary physical activity.

**Figure 7.**
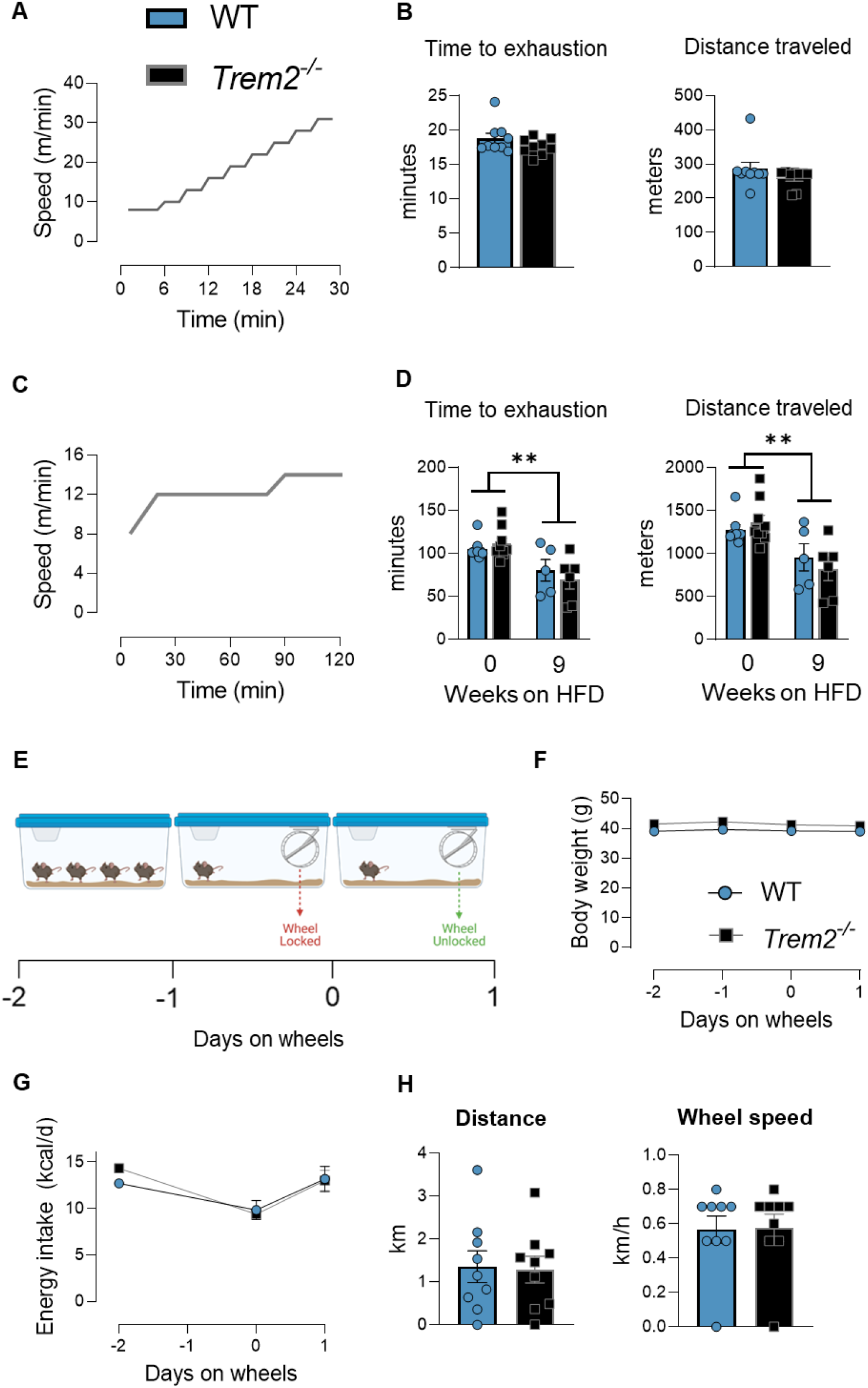
Trem2 deletion does not limit acute exercise performance. **A&B**) Peak exercise capacity and **C&D**) endurance capacity in Trem2-/- and WT mice were performed with time to exhaustion and distance traveled being the primary readout. Endurance tests were conducted in lean and obese states (9 weeks on HFD diet). **E**) Voluntary wheel running test was conducted over a 48 h period. Mice were single housed during the observation period and recombined with cage-mates after the test. Mice were transferred to a cage containing a running wheel that was physically locked from day -1 to day 0. On day 0 the wheel was unlocked and animals were allowed to run freely. **F**) Body weight, **G**) energy intake, **H**) run distance and run speed were collected. Independent t-tests were used to assess statistical differences between groups for panels B and H. Two-way ANOVA with time and genotype as factors was used to determine group differences for panel D. n=5-9/group. Trem2-/- mice were purchased directly from JAX and age-matched C57BL/6J mice were used as controls. Animal housing and experiments were conducted at room temperature (21-23°C). Graphics were generated using BioRender.com. Data are presented as mean ± SE. HFD, high fat diet. **p<0.01

## Discussion

A novel macrophage population with high expression of genes associated with lipid metabolism – termed LAMs – is enriched in adipose tissue of obese humans and rodents. LAMs are thought to be lipid-scavenging phagocytes that act as a lipid buffer during adipose tissue expansion and that subsequently spare ectopic organs from excess lipid. Trem2 is the principal defining feature of LAMs (6-8), and its expression in adipose tissue progressively increases with obesity (Fig. 1B and (6)). Trem2 deletion in mice reportedly induces AT inflammation and impairs glucose metabolism (20). Consequently, the biomedical literature is being flooded with the notion that targeting Trem2 therapeutically may be a strategy to treat metabolic diseases. Yet, convincing evidence supporting that Trem2 is a molecular switch influencing insulin resistance is lacking.

Thus, we aimed to determine whether Trem2 was causative in promoting insulin resistance in obesity. In stark contrast to previous reports, using multiple complementary approaches in male or female mice, we do not find that loss of Trem2 disrupts glucose homeostasis. Similarly, increasing metabolic demand via acute maximal or endurance exercise does not reveal disturbances in aerobic fitness nor does Trem2 deficiency perturb the acute inflammatory response to LPS. Although loss of Trem2 causes significant adipocyte remodeling and adipocyte hypertrophy, our findings cast doubt that loss of Trem2 alone is sufficient to disrupt whole-body metabolic homeostasis in lean or obese settings.

To our knowledge, four studies have determined whether Trem2 deletion or overexpression regulates glucose metabolism in mice – all of which were conducted with male animals (6, 8, 9, 20). These studies have been contrasted with the present paper in **Table 1**. The published studies implementing loss of Trem2 function report either worsened glucose tolerance (6, 8) or no effect (20). Overexpression of Trem2 also shows impaired glucose tolerance and higher fasting insulin than WT controls (9). Regardless of Trem2 loss or gain of function, three of the four papers demonstrating worsened glucose or insulin tolerance simultaneously reveal that mutant animals gain more weight than WT controls on obesogenic diets. In mice, small differences in body weight are often sufficient to delay glucose clearance following a glucose bolus. Consequently, many genotypic differences in glycemic control reported in the literature can be explained by differences in body weight or fat mass in mice. In this case, pair feeding could be used to parse out whether a genetic mutation of interest is a regulator of glucose metabolism or insulin action when body weight is matched.

Sharif et al. report that Trem2 ablation impairs glucose tolerance in weight matched obese mice (8). This is the only study, to date, to indicate a possible role of Trem2 in regulating glucose metabolism when body weight or fat mass are not different. It is important to note that these studies were performed after an overnight fast. As reported previously (13-16), overnight fasting improves glucose tolerance by enhancing insulin action in mice. It is not known if WT and Trem2-/- mice in the Sharif study lost the same percentage of body weight during the overnight fast. This is important because differences in energy stores, namely liver glycogen, at the start of the glucose tolerance test can influence hepatic glucose fluxes and by consequence the rate of blood glucose clearance. We did not directly measure glycogen stores in the present study but we did find that WT and Trem2-/- had similar reductions in body weight after a 5 h fast (WT, 3.1±0.4% vs Trem2-/- 2.9±0.2%) and 15 h fast (WT, 5.8±0.2% vs Trem2-/- 6.0±0.2%), respectively. No differences in glucose tolerance were found between genotypes in response to short or long-term fast, hence we suggest that baseline glycogen levels are likely similar between genotypes. In mouse studies, blood is routinely sampled from the tail. Sampling directly from an arterial source in an unrestrained/non-anesthetized mouse mitigates stress-induced blood glucose elevation due to elevated catecholamine’s and stress hormones (16). Thus, to avoid stress-induced hyperglycemia we surgically implanted carotid artery catheters and performed a mixed meal tolerance test via oral gavage. Consistent with other measures of metabolic control, no differences in glucose, insulin, NEFA, or triglycerides concentrations were found between genotypes. In addition, herein, mice were challenged with a thermal stressor (thermoneutrality) which is known to decrease energy expenditure and thought to decrease insulin sensitivity (21). Postprandial insulin concentrations were higher in TN versus RT housed mice, suggesting decreased insulin sensitivity, but the absence of Trem2 did not produce an interaction with housing temperature. Based on results from multiple exogenous substrates delivered, two methods of substrate administration, and two environmental conditions tested, we conclude that loss of Trem2 does not worsen glucose metabolism or insulin sensitivity in male or female mice.

**Table 1.**

| Study | Mouse model | Sex | Glucose dose (g) or kcal dose | Delivery | Fast duration | Metabolic Phenotype |  |  |  |  |
| --- | --- | --- | --- | --- | --- | --- | --- | --- | --- | --- |
|  |  |  |  |  |  | Adiposity | Fasting glucose | Glucose tolerance | Meal tolerance | Insulin tolerance |
| **Jaitin et al. (6)** | Turnball et al. (28) **Knockout**<br>Cre-Lox, homologous recombination deletion in exon 3 and 4 | M | 0.02 g (fixed dose) | i.p. | 14 h | Trem2<sup>-/-</sup> mice, elevated on HFD | Trem2<sup>-/-</sup> mice, elevated on HFD | Trem2<sup>-/-</sup> mice, impaired on HFD; no difference on LFD | ND | ND |
| **Sharif et al. (8)** | Turnball et al. (28) **Knockout**<br>Cre-Lox, homologous recombination deletion in exon 3 and 4 | M | 1 g/kg body weight | oral | overnight | Trem2<sup>-/-</sup> mice, no difference | Trem2<sup>-/-</sup> mice, no difference | Trem2<sup>-/-</sup> mice, impaired on HFD; no difference on LFD | ND | Trem2<sup>-/-</sup> mice, impaired on HFD |
| **Liu et al. (20)** | Turnball et al. (28) **Knockout**<br>Cre-Lox, homologous recombination deletion in exon 3 and 4 | M | 1.5 g/kg body weight | i.p. | 16 h | Trem2<sup>-/-</sup> mice, elevated on HFD | Trem2<sup>-/-</sup> mice, no difference | No difference on LFD or HFD | ND | Trem2<sup>-/-</sup> mice, impaired on HFD |
| **Park et al. (9)** | Park et al. (9) **Overexpression**<br>Transgene introduced via microinjection: 732 bp mouse Trem2 insertion behind pCMV promoter | M | 1 g/kg body weight | i.p. | overnight | Trem2<sup>-/-</sup> mice, elevated on HFD | Trem2<sup>-/-</sup> mice, elevated on HFD | Impaired on HFD; no difference on LFD | ND | Trem2<sup>-/-</sup> mice, impaired on HFD |
| **Present study** | JAX, #027197 **Knockout**<br>CRISPR, 175 bp deletion, stop codon in exon 2 | M&F | 2g/kg fat free mass | i.p. | 5 h and 15 h | Trem2<sup>-/-</sup> mice, no difference | Trem2<sup>-/-</sup> mice, no difference | No difference on LFD or HFD | ND | ND |
| **Present study** | JAX, #027197 **Knockout**<br>CRISPR, 175 bp deletion, stop codon in exon 2 | M&F | 3 g/kg fat free mass | oral | 5 h | Trem2<sup>-/-</sup> mice, no difference | Trem2<sup>-/-</sup> mice, no difference | No difference on HFD or LFD | ND | ND |
| **Present study** | JAX, #027197 **Knockout**<br>CRISPR, 175 bp deletion, stop codon in exon 2 | M&F | 3.5 g/kg fat free mass | oral | 5 h | Trem2<sup>-/-</sup> mice, no difference | Trem2<sup>-/-</sup> mice, no difference | No difference at standard housing temperature or thermoneutrality | ND | ND |
| **Present study** | JAX, #027197<br>**Knockout**<br>CRISPR, 175 bp deletion, stop codon in exon 2 | M&F | 12 kcal/kg body weight | oral | 5 h | Trem2<sup>-/-</sup> mice, no difference | Trem2<sup>-/-</sup> mice, no difference | NA | No difference on HFD | ND |

ND, not determined NA, not applicable M, males F, females

Acute exercise exerts a major influence on energy metabolism that requires a coordinated response to mobilize and utilized substrates for mechanical work. Given the large adipocyte size in Trem2-/- mice we hypothesized that adipocyte-derived NEFA via lipolysis may be impaired when mice are challenged with exercise. If this hypothesis were correct, mice would be forced to oxidize glucose as the major fuel source and fatigue more quickly than WT animals.

However, Trem2 deficient animals did not manifest differences in peak graded exercise performance or endurance exercise (i.e., a metabolic state in which lipolysis is high and sustained) in either lean or obese states. We acknowledge that these exercise tests are limited by ‘time to exhaustion’ being the primary readout. It is plausible that substrate utilization (respiratory quotient) is different between genotypes. Higher reliance on carbohydrate as a fuel source is closely linked to exercise time to exhaustion during an endurance-based test (22, 23); therefore, we postulate that if the respiratory quotient was indeed affected by loss of Trem2 this would have manifested in time to exhaustion. Future experiments would benefit from indirect calorimetry to determine maximal oxygen consumption and respiratory exchange ratio’s (e.g. estimate of whole-body substrate utilization) during exercise tests.

Although Trem2 does not appear necessary for acute regulation of exercise responses, this does not rule out a role in chronic exercise adaptations. In rodent Alzheimer’s disease models, the anti-inflammatory effects of exercise training is associated with Trem2 expression levels (10, 24). In addition, exercise training blunts the increase in soluble Trem2 caused by neuroinflammation in mice (24), but not in human patients with Alzheimer’s disease (25). Nonetheless, given that metabolic function (i.e., glucose and lipid metabolism) is not adversely affected by loss of Trem2 in the present study, it seems unlikely that peripheral metabolic adaptations to exercise training would require Trem2. In addition, in the absence of disease (controls used in rodent Alzheimer’s models (10, 24)), exercise does not further enhance Trem2 expression, suggesting that exercise may prevent the suppression of Trem2 expression following microglia activation, but exercise does not seem to increase Trem2 expression above basal. It may be that exercise motivation or recovery to muscle damage (i.e., mild injury) are influenced by Trem2 expression. Additional research is needed to determine whether the role of Trem2 in tissue remodeling is linked with exercise training-induced adaptations.

In response to energy excess, Trem2 appears essential for the beneficial functions of LAMs, including lower inflammation and by placing a break on adipocyte hypertrophy (6). Some studies report enhanced inflammatory gene expression in adipose tissue isolated from Trem2-/- mice (20), whereas we did not observe worsened overall adipose tissue inflammation in obese Trem2-/- mice (Fig. S2A) or when challenged acutely with a bacterial pathogen (Fig. 2). The abundance of crown-like structures, but not total macrophages, in adipose tissue is lower in Trem2-/- versus WT animals, which could result in delayed clearance of dying adipocytes, and by consequence, increased inflammation in some studies. Despite using a different model of Trem2 deletion herein, the adipose tissue phenotype is similar to published literature with Trem2-/- having excessive adipocyte hypertrophy and dysregulated expression of LAM transcriptional profile. In response to a lethal dose of endotoxin, prior data reveal that obese Trem2-/- mice have accelerated death rates, which was attributable to an energetic crisis in the liver (12). *Trem2* expression is suppressed in response to LPS and does not increase until 48-72 hours after endotoxin *in vivo* (26) and *in vitro* (Fig. 2I) – well after the inflammatory phase.

This suggests Trem2 may be involved in inflammatory recovery or reestablishing homeostasis. A mild delay in plasma cytokine clearance (15-18h post LPS) has been documented in mice with a Trem2 missense mutation or global knockout (26, 27). Yet, this delay in cytokine clearance manifests before Trem2 expression increases, thus causation is still unclear.

Although *in vitro* data imply an important role for Trem2 in inflammatory recovery, whether these observations translate *in vivo* requires additional study. Evidence from the latter may expand our understanding of the relationship between macrophage lipid handling and tissue injury.

In conclusion, we report a disassociation between adipose tissue remodeling caused by loss of Trem2 and whole-body metabolic homeostasis in mice. We reject the hypothesis that Trem2 is a molecular trigger underpinning insulin sensitivity and rather suggest that Trem2 may be important for the resolution of inflammatory perturbations and tissue remodeling that accompany weight gain.

## ACKNOWLEDGEMNTS

We acknowledge the following Vanderbilt University (VU) and Vanderbilt University Medical Center (VUMC) core facilities: VUMC Hormone Assay & Analytical Services Core (NIH DK059637 and DK020593), VU Metabolic Mouse Phenotyping Center [VMMPC (NIH DK059637; www.vmmpc.org)], and Translational Pathology Shared Resource (NCI/NIH Cancer Center Support Grant 5P30 CA68485-19). We thank Alec Rodriguez (VU), Vitrag Patel (VU), and Amber Crabtree (VU) for mouse husbandry and technical assistance.

## AUTHOR CONTRIBUTIONS

NCW conceived and designed research, performed experiments, interpreted results, and drafted the manuscript. EMW performed experiments, interpreted results, and reviewed/edited the manuscript. JNG performed experiments, interpreted results, and reviewed/edited the manuscript. AHH conceived and designed research, interpreted results, and reviewed/edited the manuscript. All authors approved the final version of the manuscript. The guarantor for this work is Dr. Alyssa Hasty.

