## Supplemental Data for "Trem2 deficiency does not worsen metabolic function in diet-induced obese mice"

**Supplemental Figure 1**

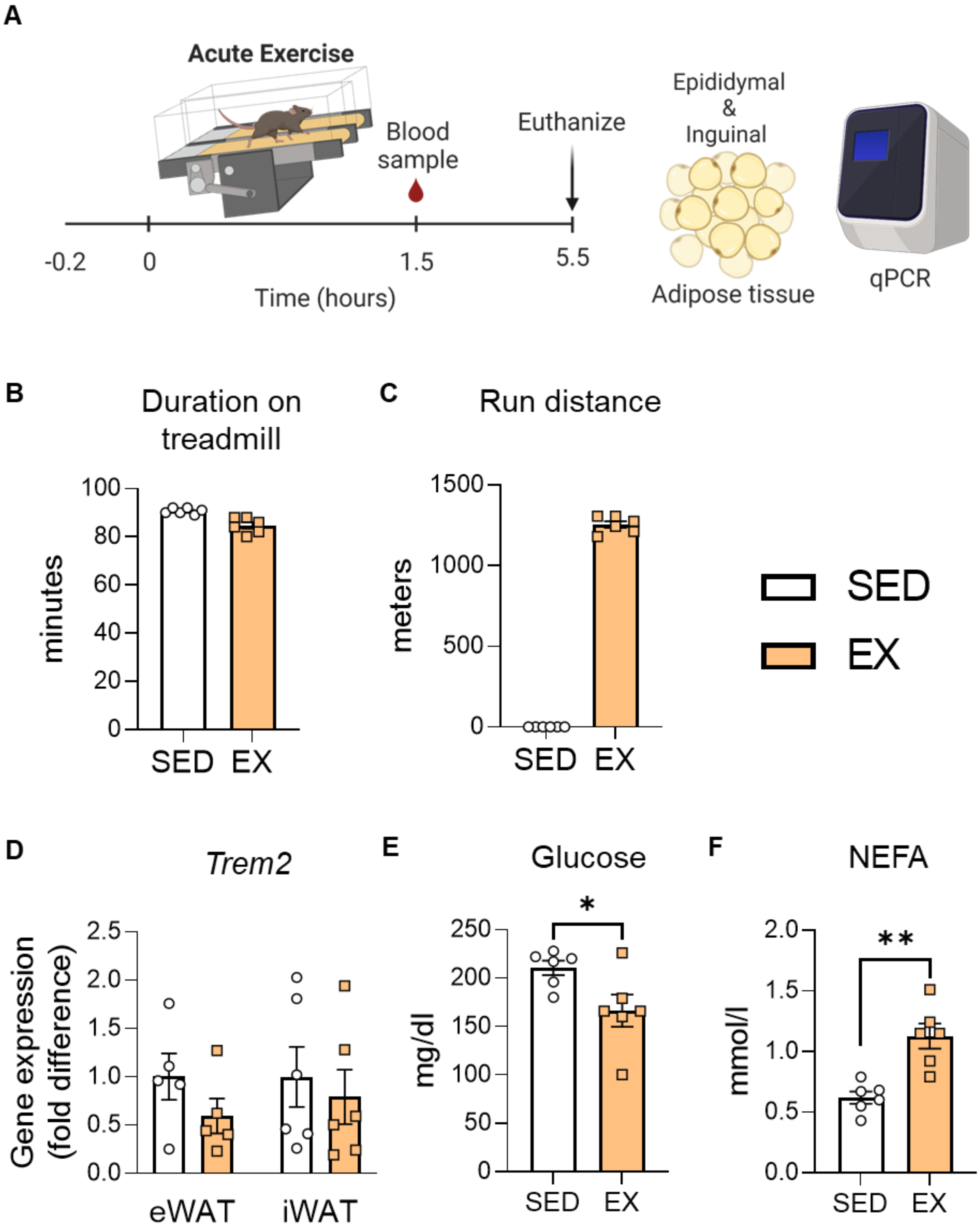

**Supplemental Figure 1** – *Acute exercise does not modify Trem2 expression in adipose tissue*. A) Acute exercise challenge was performed in lean male WT and Trem2-/- mice. B) Mice habituated to a motorized treadmill ran at a moderate intensity for approximately 90 minutes, while sedentary mice remained on the treadmill belt in the OFF setting. C) By design the run distance was zero in the sedentary group and was approximately 1200 meters in the exercise group. A blood sample was obtained immediately after the exercise bout to confirm known exercise responses of glucose and NEFA. Four hours after the exercise bout, mice were euthanized and adipose tissue depots were collected and flash frozen. D) mRNA expression for Trem2 was assessed in epididymal and inguinal adipose tissue. No differences in Trem2 expression were found between sedentary and acute exercised mice. E) Glucose was decreased immediately following exercise whereas F) NEFA concentrations were increased, consistent with the known responses to acute exercise. These data suggest that adipose tissue Trem2 is not responsive to acute exercise (4 h post exercise; n=6/group). Trem2-/- mice were purchased directly from JAX and age-matched C57BL/6J mice were used as controls. Independent *t* tests were used to assess statistical differences between groups. Graphics were generated using BioRender.com. Data are presented as mean ± SEM. * $p$<0.05, ** $p$<0.01. SED, sedentary; EX, exercise; NEFA, non-esterified fatty acids; eWAT, epididymal adipose tissue; iWAT, inguinal adipose tissue.

**Supplemental Figure 2**

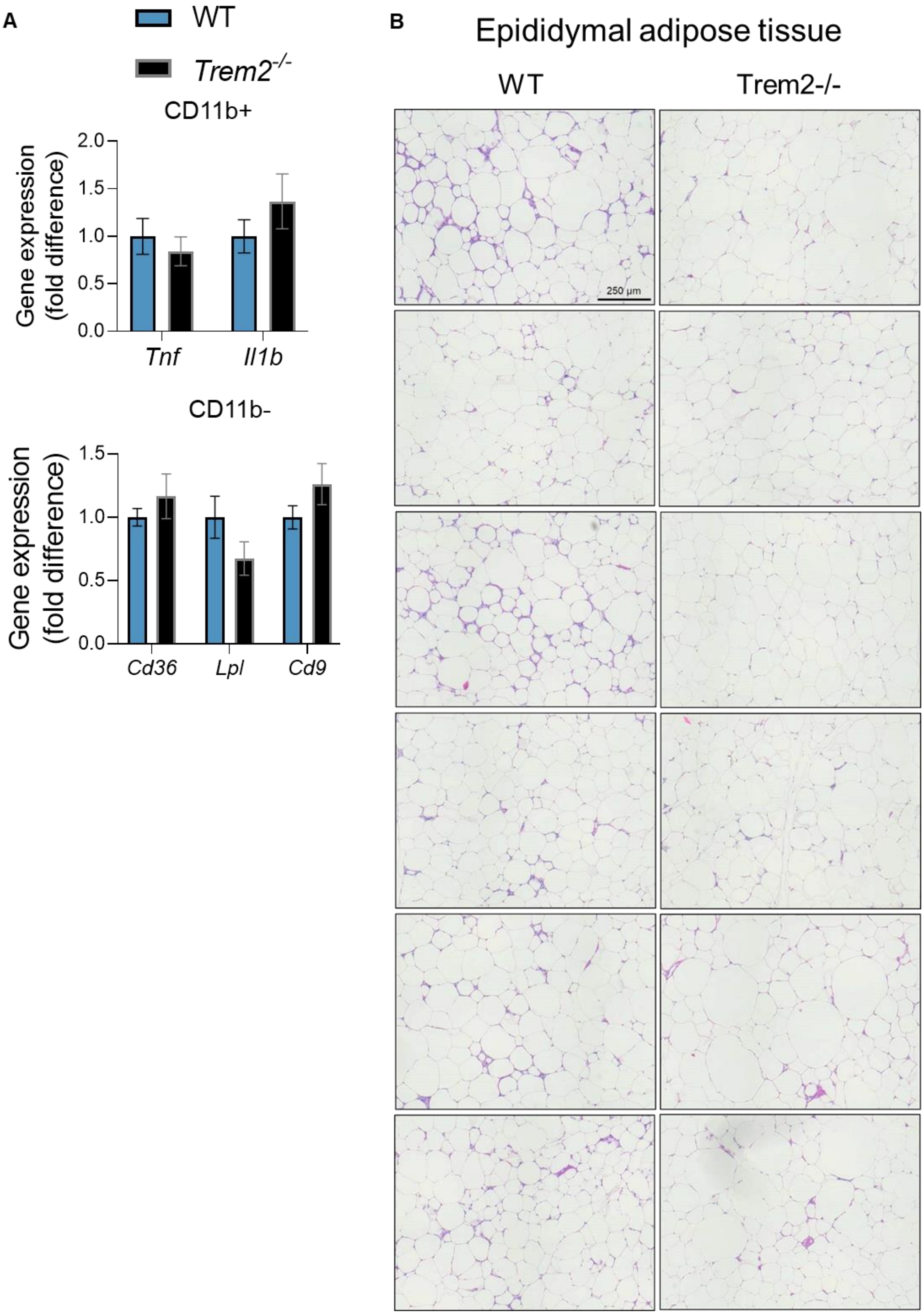

**Supplemental Figure 2** – *Loss of Trem2 does not potentiate adipose tissue inflammatory mRNA expression*. Male mice were fed high fat diet for 18 weeks. Epididymal adipose tissue was extracted and sorted for CD11b+ and CD11b- fractions. A) Enriched cell populations were prepared for RNA extraction and cDNA synthesis. mRNA expression for *Tnf* and *Il1b* were determined in CD11b+ cells, which did not reveal differences between genotypes (n=7-8/genotype). CD11b- fraction was assessed to lipid associated macrophage (LAM) markers to determine whether Trem2 affected non-macrophage populations. Non-macrophage enriched cells did not manifest differences in LAM markers suggesting that Trem2 was specific to myeloid cells (n=7-8/genotype). B) Individual adipose tissue H&E sections are presented from 18-week high fat fed male WT versus Trem2-/- mice. Overall, it is clear that 1) WT mice appear to exhibit more crown-like structures as indicated via deep purple staining surrounding individual adipocytes and 2) Trem2-/- sections are interspersed with excessively large adipocytes. Independent *t* tests were used to assess statistical differences between groups. Data are presented as mean ± SEM.
